# Wing structure and neural encoding jointly determine sensing strategies in insect flight

**DOI:** 10.1101/2021.02.09.430476

**Authors:** Alison I. Weber, Thomas L. Daniel, Bingni W. Brunton

## Abstract

Animals rely on sensory feedback to generate accurate, reliable movements. In many flying insects, strain-sensitive neurons on the wings provide rapid feedback that enables stable flight control. While the impacts of wing structure on aerodynamic performance have been widely studied, the impacts of wing structure on sensing remain unexplored. In this paper, we show how the structural properties of the wing and encoding by mechanosensory neurons interact to jointly determine optimal sensing strategies and performance. Specifically, we examine how neural sensors can be placed effectively over a flapping wing to detect body rotation about different axes, using a computational wing model with varying flexural stiffness inspired by the hawkmoth *Manduca sexta*. A small set of mechanosensors, conveying strain information at key locations with a single action potential per wingbeat, permit accurate detection of body rotation. Optimal sensor locations are concentrated at either the wing base or the wing tip, and they transition sharply as a function of both wing stiffness and neural threshold. Moreover, the sensing strategy and performance is robust to both external disturbances and sensor loss. Typically, only five sensors are needed to achieve near-peak accuracy, with a single sensor often providing accuracy well above chance. Our results show that small-amplitude, dynamic signals can be extracted efficiently with spatially and temporally sparse sensors in the context of flight. The demonstrated interaction of wing structure and neural encoding properties points to the importance of their joint evolution.

## Introduction

The physical structure of an animal’s body transforms the incoming sensory information and can either facilitate or constrain sensing capacity. Indeed, body parts in many systems serve to preprocess sensory inputs in ways that are beneficial for the organism, extracting relevant features and reducing downstream computational burdens [1]. For instance, the decrease in stiffness from base to apex of the mammalian cochlea promotes frequency selectivity along its length [2], and the response properties of mechanosensors that encode high-frequency vibrations in mammalian skin are determined largely by the viscoelastic properties of layered, fluid-filled capsule surrounding nerve endings [3]. On the other hand, structure may limit sensing by imposing constraints on the numbers or locations of sensory receptors. In the vibrissal system, for example, mechanosensory neurons are located only at the whisker base, so information about an object’s point of contact with the whisker cannot be directly measured [4]. Thus, the physical properties of non-neural structures play an important role in determining how stimuli are experienced and transduced by sensory receptors embedded in the body.

Sensory receptors transduce stimuli into electrical signals, typically taking the form of action potentials, in which the incoming signal is converted into a series of all-or-none events. Sensory neurons respond selectively to particular features of the stimulus; for instance, auditory neurons in the cochlea respond to particular frequencies, and visual neurons in the retina respond to distinct temporal patterns of illumination [5, 6, 7]. This transformation from input to response can often be accurately represented by a model of neural encoding that characterizes the stimulus features to which the neuron is sensitive along with a nonlinear function that captures how selective the neuron is for those features [8, 9]. Importantly, neural encoding properties define the information available to the rest of the nervous system about the animal’s environment and its own body. These properties may adapt across multiple timescales, changing over rapid timescales based on the history of inputs as well as over evolutionary time.

Here we focus on the interaction of body structure and sensory encoding properties in the context of insect flight control, where wings provide rapid sensory feedback necessary for stable flight [10, 11, 12, 13, 14]. Although extensive previous work has examined how wing structure impacts aerodynamic performance [15, 16, 17, 18, 19, 20], the impacts of wing structure on sensing remain unexplored. Structural properties (e.g., geometry, flexural stiffness) interact with forces acting on the wing to produce local spatiotemporal patterns of strain that are sensed by the nervous system. While forces are dominated by wing flapping, body rotations elicit small Coriolis forces in the wings that are several orders of magnitude smaller, making the detection of body rotation a challenging sensing task. Local strain is encoded by sensors, called campaniform sensilla, distributed sparsely over the wings at consistent locations across individuals. It has recently been demonstrated that only a small number of sensors are needed to read out behaviorally relevant information about body rotations, with neural sensors providing advantages over sensors that directly encode strain [21]. Sparse sensing strategies have advantages in terms of both energetic cost and robustness [22, 23, 24, 25]. However, no previous work has examined the impacts of wing structure on sparse sensing strategies.

In this paper, we show that just a few spiking sensors can detect subtle differences in strain that arise from the relatively tiny Coriolis forces produced during combined wing flapping and body rotation. We use a simple model based on *Manduca sexta* to characterize spatiotemporal patterns of strain over the wing during flapping. We then encode this strain in a population of spiking sensors, which we refer to as neural-inspired sensors, whose response properties are based on experimental measurements from mechanosensory neurons. With a sparsity-promoting optimization method, we solve for the locations of a small, fixed number of sensors for detecting body rotation about different axes. We focus on wing stiffness because it is an important structural property that varies widely across species [26], within species [27], and even over an animal’s lifetime [28]. We show that wing stiffness and neural encoding properties jointly determine the accuracy of body rotation detection and optimal sensor locations. Moreover, sensing performance is remarkably resilient to sensor loss and external disturbances, indicating that sparse sensor placement can be used for efficient, robust sensing in the context of flight.

## Results

To understand how sensing strategies and performance are impacted by wing structure, we focused on one specific sensing goal: finding a minimal set of strain-encoding, spiking mechanosensors on the wing that are effective in detecting body rotation. Given the timing of a single spike from each of a set of sensors placed at optimal locations, we ask how well rotation of the body about different axes can be discriminated. We then analyze this performance and the optimal locations of the sensors on the wings, assessing how they change as we manipulate wing stiffness and neural encoding properties.

We first simulate spatiotemporal patterns of strain over a wing using an Euler-Lagrange model inspired by the wings of the hawkmoth *Manduca sexta* [29] (Fig 1A, top). Strain data are acquired for two conditions: one in which the wing is flapping, and another in which the body is undergoing rotation while the wing is flapping. The combination of flapping and rotation leads to Coriolis forces that alter the spatiotemporal patterns of strain, exciting small torsional modes that are absent when flapping alone [29]. The time history of strain is then encoded by a population of neural-inspired sensors, whose encoding properties are based on experimental measurements of mechanosensors in hawkmoth wings [12]. The output of this neural encoding step is a pattern of spikes, temporally sparse all-or-none signals (Fig 1A, middle). We then solve for a sparse set of sensors that can be used to detect body rotation based on spike timing (Fig 1A, bottom). A single spike time from each sensor in this subpopulation is used to determine whether or not the wing is undergoing rotation at each wingbeat. We carry out this procedure for wings of different stiffness (Fig 1B, top) and sensors with different thresholds (Fig 1B, top) to assess how sensing strategies depend on wing structure.

**Fig 1.**
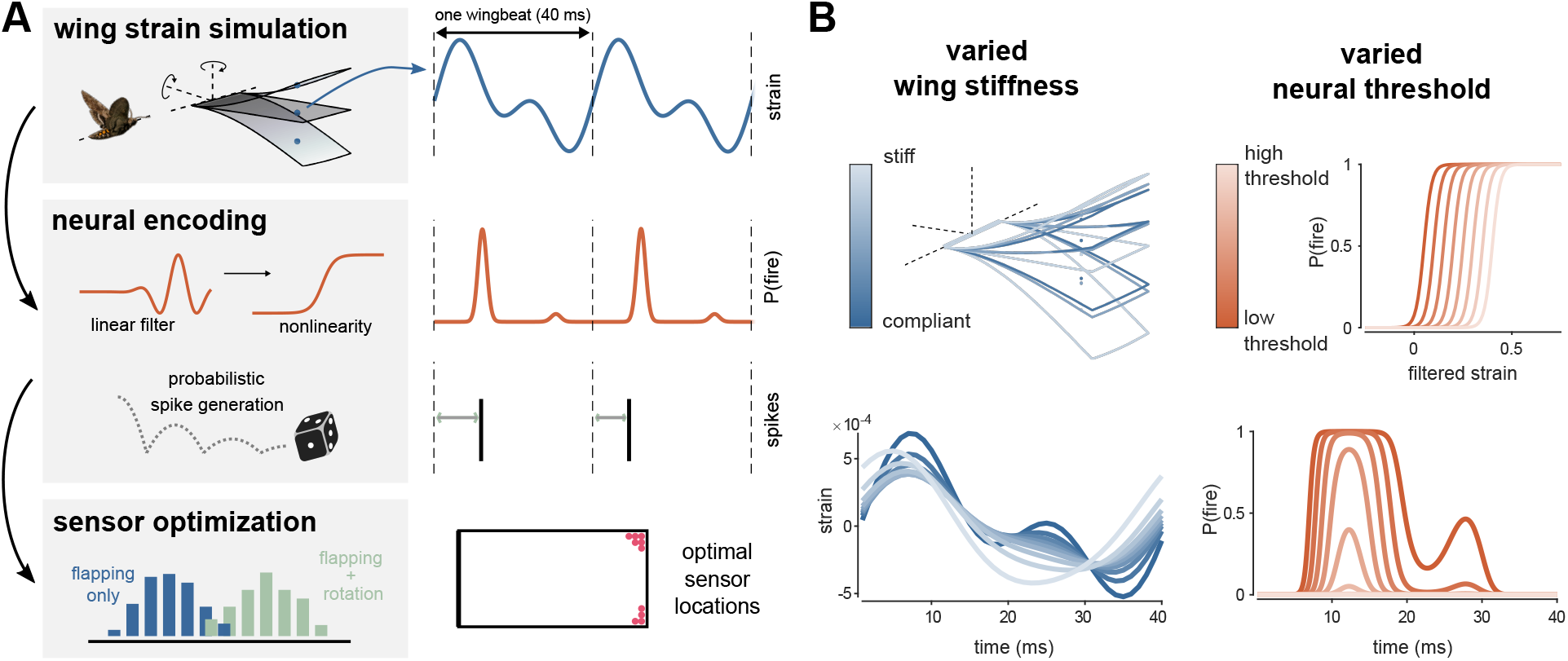
Wing stiffness and neural encoding properties determine optimal sensor locations on a flapping wing. A, *Top:* Strain over time is simulated at each possible sensor location in an Euler-Lagrange model of a flapping wing [29, 21]. The wing may be subject to rotation in different axes. Strain is shown for an example sensor location (blue dot). *Middle:* Strain at each location is encoded by a neural-inspired sensor. Strain is convolved with a linear filter and passed through a static nonlinearity to generate a probability of firing a spike over time, P(fire). Spikes are probabilistically generated, so that exact spike time varies from wingbeat to wingbeat. *Bottom:* The full dataset, a matrix of spike times at all sensor locations for each wingbeat, is used to find a subset of sensors from which information about rotation can be read out. Linear discriminant analysis is used to find the vector *w*, such that when the data are projected onto *w* the means of the flapping only and flapping with rotation data are maximally separated. B: *Left:* Changing wing stiffness produces different wing movements (top, shown at 3 time points for 3 different stiffness values) and different strain (bottom, shown for identical location, indicated by dot, on wings of different stiffness) over the course of each wingbeat. *Right:* Changing the threshold of the neural encoding nonlinearity (top) alters of the probability of firing over the course of the wingbeat (bottom), with higher thresholds resulting in lower probability.

### Spike timing and precision determine local peaks in accuracy as a function of wing stiffness

We vary wing stiffness while holding neural encoding properties constant (linear filter frequency parameter *ω* = 1; neural threshold *β* = 0.2). We find sparse sensor locations that can detect rotation in the yaw axis and evaluate performance using the 10 best sensors. Classification accuracy changes nonmonotonically with wing stiffness, peaking near 100% accuracy at a wing stiffness similar to that for hawkmoth wings (3 GPa, stiffness factor = 1). There is a second, smaller peak near 75% at a much lower stiffness (Fig 2A). In both cases, the optimal sensor locations occur at the wing tip. The single best sensor location for each of these stiffness values is shown in Fig 2A, although all of the 10 best sensors in each case fall at the wing tip and show similar patterns of strain and spiking over time.

**Fig 2.**
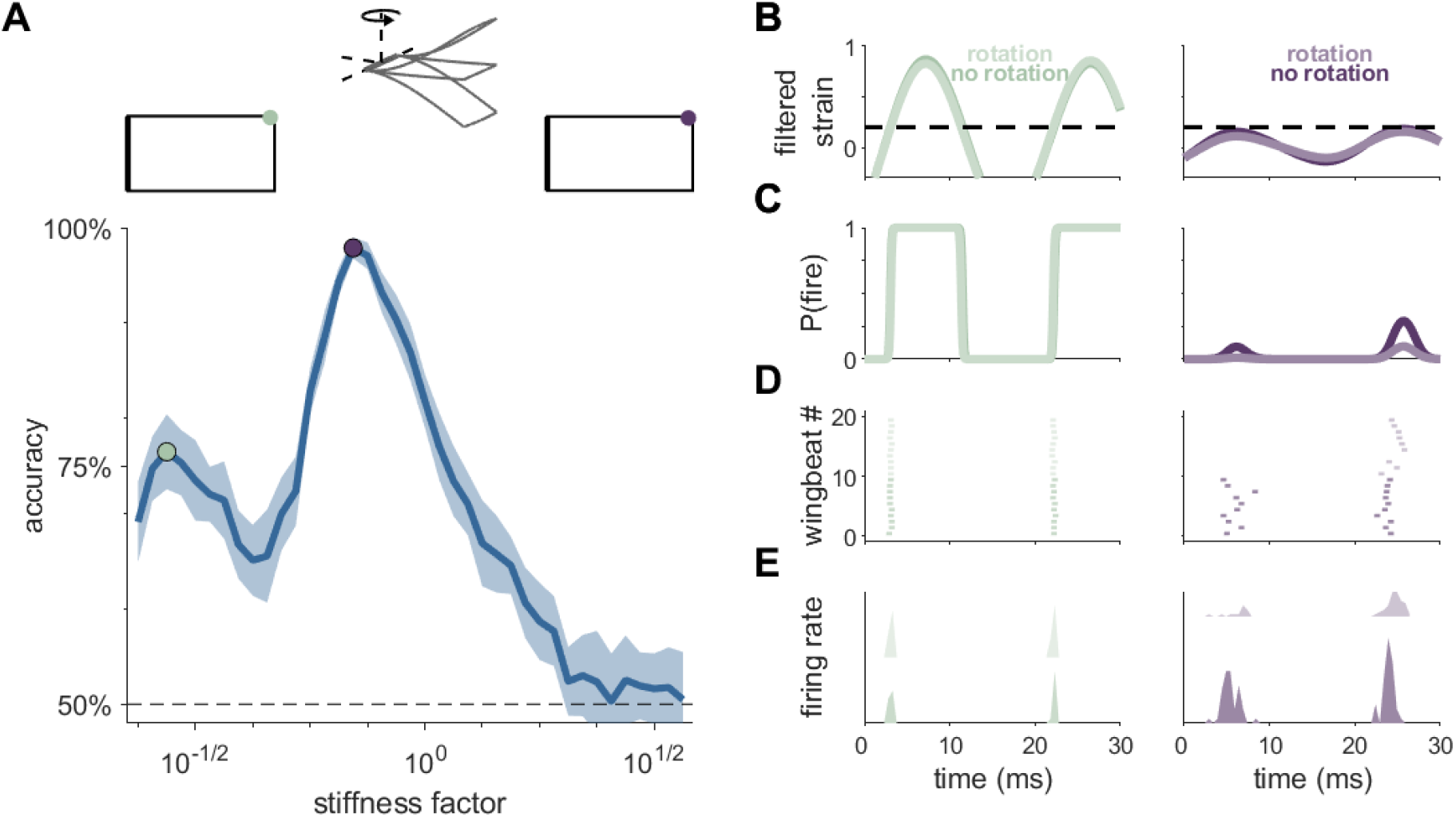
Wing stiffness interacts with neural threshold to determine spike timing. A: Accuracy as a function of wing stiffness. Chance accuracy is 50%, and stiffness factor 1 corresponds to 3 GPa, comparable to experimentally measured stiffness in the hawkmoth [26]. Green and purple dots indicate local peaks in accuracy. Shaded area indicates ±1 standard deviation. *Top:* Center schematic indicates the direction of rotation. Left and right schematics show the location of the single best sensor for each stiffness value. B: Filtered strain over a single wingbeat for flapping only (“no rotation”) and flapping with rotation (“rotation”) conditions at a single sensor location. Green and purple dots in A correspond to left/green and right/purple filtered strain. Dashed horizontal line indicates the threshold (value at half-max) of the neural encoding nonlinearity. C: Filtered strain passed through the nonlinear function gives probability of spiking over time, P(fire). D: Spiking responses for 10 wingbeats each of the no rotation and rotation conditions. Spikes are generated probabilistically from P(fire), resulting in variable spike timing from wingbeat to wingbeat. E: Histogram of spike times (PSTHs) for each condition, summarizing spike timing over hundreds of wingbeats.

Differences in spike timing and spike timing precision underlie these local peaks in accuracy. For more compliant wings, the filtered strain in optimal sensors quickly crosses threshold (Fig 2B, left), leading to sharp transitions from zero probability of firing (P(fire)) a spike to certainty of firing a spike (P(fire) ~100%, Fig 2C, left). This results in precisely timed spikes both when the wing is flapping only and flapping with rotation (Fig 2D,E, left). Small differences in filtered strain under these two conditions translates to small but detectable differences in spike timing. With 10 sensors, wing rotation can be detected with ~75% accuracy.

For stiffer wings, the filtered strain in optimal sensors varies less over time, barely reaching threshold (Fig 2B, right). This results in an overall lower probability of firing but also amplifies the difference between flapping only and flapping with rotation: small differences in filtered strain translate to clear differences in P(fire) over time (Fig 2C, right). This is reflected in spike timing, with the first spike being much more likely to occur earlier in the wingbeat cycle in the flapping case (around 5 ms) compared to flapping with rotation (around 25 ms, Fig 2D,E, right). Lower precision in spike timing is offset by large differences in the time to the first spike, resulting in a classification accuracy approaching 100%.

### Structure and neural encoding interact to determine accuracy and optimal sensor locations

To explore how neural encoding properties interact with wing stiffness, we varied the threshold of the nonlinearity in neural encoding. The threshold determines how selective the sensor will be for the feature given by the linear filter, with higher thresholds imparting stronger selectivity. The slope of the nonlinearity most strongly affects spike timing precision, with higher slopes leading to faster transitions from P(fire) = 0 to 1 and therefore greater spike timing precision. We set the slope of the nonlinearity to roughly match the spike timing precision observed in wing mechanosensors (0.1-1 ms, [11]). The frequency content of the linear filter was chosen based on experimental observations [12]. Higher-frequency filters show uniformly low performance as a function of wing stiffness, while lower frequency filters show uniformly high performance (Supp Fig 2).

The ability to detect body rotations depends jointly on wing stiffness and neural threshold, with very dissimilar combinations yielding similarly high classification accuracy (Fig 3A). For example, for stiffness factors just below 1, accuracy is similarly high for low (0.15) and high (0.7) thresholds, but accuracy is lower at intermediate values (e.g., 0.3).

**Fig 3.**
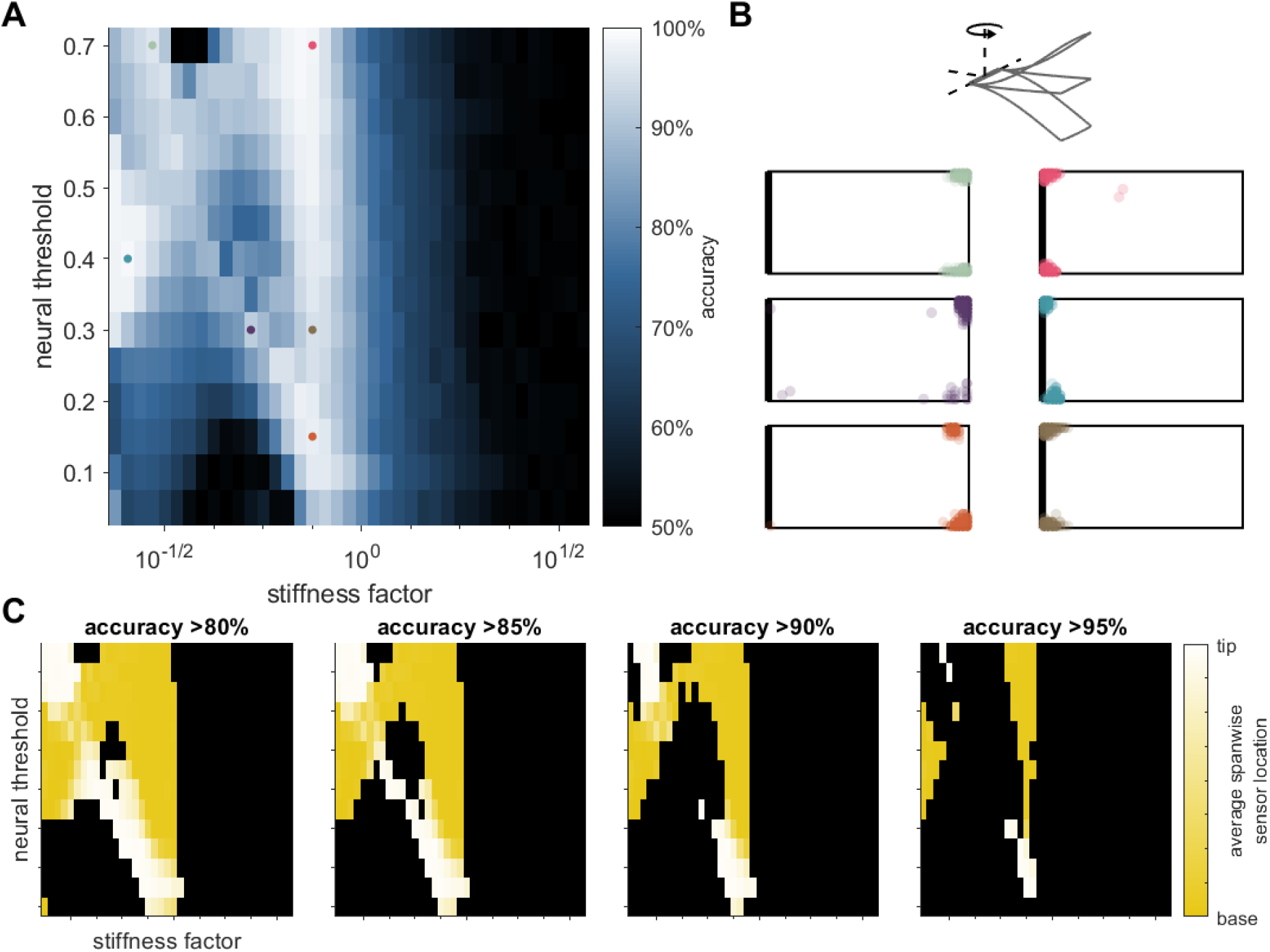
Wing stiffness and neural threshold interact to determine sensor locations and classification accuracy in the yaw axis. A: Accuracy as a function of neural threshold and wing stiffness. B: Optimal sensor locations of 10 best sensors, overlaid for 20 different simulated datasets. Neural threshold and wing stiffness of each panel are indicated by the corresponding colored dot in A. Top schematic indicates the axis of rotation. C: Optimal sensor locations in the spanwise direction from wing base (yellow) to wing tip (white) as a function of neural threshold and wing stiffness. Each panel shows results for a different accuracy cutoff. Black indicates parameter combinations that fall below the cutoff.

Moreover, the optimal sensing strategies for a given wing stiffness can be qualitatively different for high and low thresholds. At stiffness factors just below 1, optimal sensors are located at the corners of the wing tip for low thresholds (Fig 3B, orange) and at the corners of the wing base for high thresholds (Fig 3B, pink). Note that sensors are expected to fall at the wing corners, as observed in previous work [21], because rotation elicits a twisting mode in wing bending, which leads to the largest differences in strain at wing corners between rotating and non-rotating conditions [29]. However, the distinction between sensors located at the wing base and those at the wing tip was not seen in previous work and is not necessarily expected [21]. When the spanwise sensor location (wing base to wing tip) is plotted as a function of neural threshold and wing stiffness, distinct regions of qualitatively different encoding strategies emerge (Fig 3C). Interestingly, these regions have rather convoluted, discontinuous shapes, illustrating that the optimal sensing strategy reflects the complex interaction of structural properties and neural encoding properties.

### For a different classification task, a simple encoding principle emerges

We next perform a similar set of numerical experiments for detecting rotation in the pitch axis. Classification accuracy for this task is generally lower than for detecting rotation in the yaw axis, with maximum accuracy across all parameter combinations tested only 77% (Fig 4A). Several clear differences emerge between the classification of pitch and yaw. First, classification accuracy for pitch is near chance for all but the stiffest wings tested, whereas the accuracy for yaw is near chance only for the stiffest wings. Second, optimal sensors are located near the wing base for all parameters tested (Fig 4B). Unlike the classification of yaw, there is a straightforward relationship between wing stiffness and accuracy, with little dependence on neural threshold. A single best encoding strategy emerges, with optimal sensors at the wing base.

**Fig 4.**
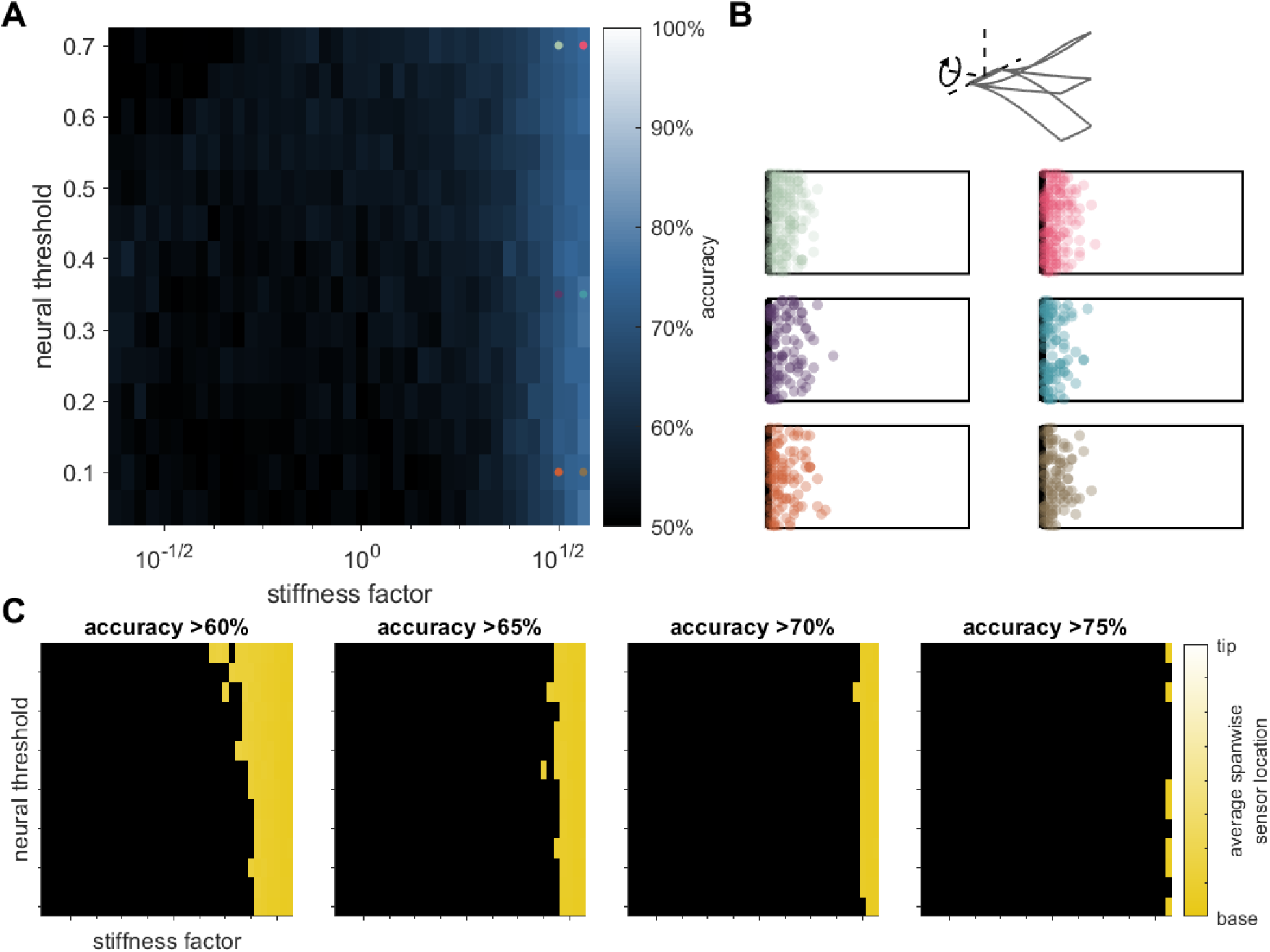
For detecting rotation in the pitch axis, sensor locations do not depend on stiffness or neural encoding properties. A: Accuracy as a function of neural threshold and wing stiffness. B: Optimal sensor locations of 10 best sensors, overlaid for 20 different datasets. Neural threshold and wing stiffness of each panel are indicated by the corresponding colored dot in A. Top schematic indicates the axis of rotation. C: Optimal sensor locations in the spanwise direction from wing base (yellow) to wing tip (white) as a function of neural threshold and wing stiffness. Each panel shows results for a different accuracy cutoff. Black indicates parameter combinations that fall below the cutoff.

### Robustness to perturbations

To determine the extent to which our results are robust to different perturbations, we test two types of perturbations that reflect likely disruptions to both biological and engineered systems: external disturbances and sensor loss.

Sensor loss may arise from wing damage sustained over an animal’s lifetime or due to sensor failure in engineered systems. We select two wing stiffness/neural threshold parameter combinations that yield near-peak accuracy ( ~100% for yaw rotation and ~75% for pitch rotation). For yaw axis rotation, one parameter set results in sensors placed at the wing base and one at the wing tip; for pitch axis rotation, both are located at the wing base. We randomly eliminate between 1 and 9 out of 10 sensors and compute accuracy using the remaining sensors. Accuracy falls smoothly as sensors are randomly dropped with, surprisingly, a single sensor still providing ~75% accuracy on average for yaw and ~60% accuracy for pitch (Fig 5A,C). Accuracy falls more gradually for pitch detection, likely due to the lower starting accuracy.

**Fig 5.**
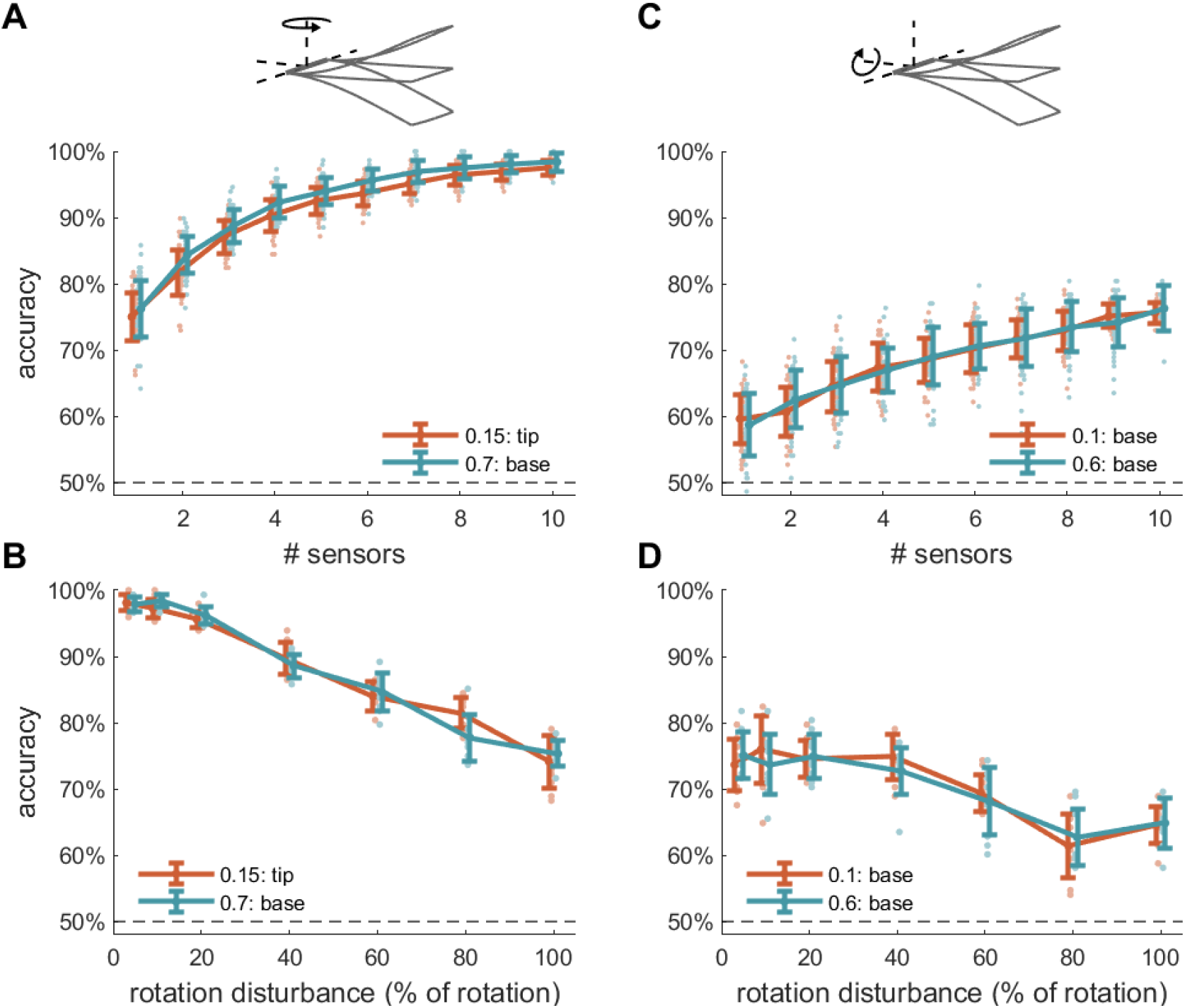
Robustness to sensor dropout and external disturbance. A: Accuracy of detecting rotation in the yaw axis as a function of number of sensors for two different wing stiffness/neural threshold parameter combinations (orange and teal). One parameter combination results in sensors at the wing tip, and a second results in sensors at the wing base, both with accuracy of 98% for 10 sensors. B: Accuracy as a function of the standard deviation of rotation disturbance, reported as the percentage of the constant rotation velocity to be detected. Parameter combinations same as A. C,D: Same as A,B for rotation in pitch axis. Two different parameter combinations were again chosen that produce similar accuracy with differing neural thresholds. For pitch-axis rotation detection, all sensors are located at the wing base. Error bars in all panels denote ±1 standard deviation.

We next examine the effects of external disturbances to wing rotation, which may arise in either biological or engineered systems due to environmental fluctuations such as wind gusts. For both yaw and pitch rotation, accuracy falls gradually as a function of the disturbance magnitude, with accuracy still well above chance when the standard deviation of the disturbance is the same magnitude as the rotation to be detected (Fig 5B,D).

In summary, for both types of perturbation, accuracy falls smoothly as the perturbation increases, with no indication of catastrophic failure at some threshold level of perturbation. Moreover, the trends are nearly identical for very different sensors, even when those sensors are located on opposite ends of the wing and have very different neural thresholds. We therefore see no evidence of differences in robustness between sensors at different locations or with different thresholds.

## Discussion

Our work is a first step toward understanding how insect wing structure determines incoming sensory information and thus sensing strategies. We examine the impacts of wing structure, namely stiffness, on sensory encoding in a computational model of an insect wing. We show that a small number of sensors, conveying a spatially and temporally sparse signal, can be used to reliably detect rotation over a range of wing stiffness values. Optimal encoding properties of these sensors and their placement vary with both wing stiffness and the need to identify rotation in different axes. Sensors are clustered at either the wing base or wing tip, depending on a combination of these factors. Moreover, accuracy is robust to multiple types of perturbations, and robustness does not depend on the particular details of neural encoding or sensor placement. The interaction of wing structure and neural encoding properties points to the importance of considering how these features may evolve together to enable sensing.

### Body structure transforms sensory inputs

An animal’s body transforms incoming sensory information in a variety of ways prior to its transduction by sensory neurons, such as the pupil and lens prior to its arrival at the retina or the mechanics of skin affecting how deformations are transmitted to embedded sensory structures [30, 31]. In systems that rely on mechanosensation in particular, there is a large body of work pointing to the importance of structure in preprocessing sensory inputs (reviewed in [1]). Mechanical properties of a structure filter an incoming signal, such as the middle ear in audition or skin in touch preferentially transmitting certain frequencies [32, 31]. Resonance can also amplify certain frequencies, such as in hair cells and rodent whiskers [33, 34, 35]. In insect flight control, considerable attention has focused on the mechanical responses of halteres, dipteran mechanosensory organs derived from hindwings [36, 37, 38, 39]. In contrast, insect wings are rarely considered in their role as sensory structures.

Future work using wing models based on the finite elements method (FEM) would allow for greater flexibility in several aspects of wing strain simulation than the Euler-Lagrange model used in the present work. More realistic wing geometry, wing venation patterns, and non-uniform stiffness all present interesting questions that can be tackled with FEM [27, 40]. The consequences of many of these properties have been previously explored in a variety of models, although none from a sensing perspective [27, 17, 16, 41, 42].

In addition to the passive transformation of sensory inputs performed by body structure, animals also engage in active sensing strategies, in which the animal’s behavior further impacts the incoming information [43, 44, 45]. Rodents, for example, modifying whisking frequency based on external conditions [46]. Behavioral changes as subtle as posture can also impact the frequency content of vibrations transmitted through the body [47]. Future work in insect wings could similarly consider the trajectory of the wingstroke as a mechanism for active sensing.

### Sensing for flight in engineered and biological systems

Developing sensing methods – including sensor number, placement, response properties, and sensor readout – has long been of interest to engineers who seek to build efficient, lightweight, low-cost systems. Most work in the domain of flight has focused on sensing in fixed-wing aircraft, rather than in flapping flight as exhibited by flying animals [48, 49, 50, 51, 52]. Similar principles of optimality, efficient sensing, sparsity, and cost reduction have also informed neuroscientists’ understanding of the nervous system [53, 54, 55, 24, 56]. In many cases, the encoding properties of sensory neurons are matched to the statistics of the incoming signals. These encoding properties may be tuned over evolutionary time, or dynamically on subsecond timescales to match changing statistics in the environment [57, 58]. Understanding optimal sensing strategies therefore provides a useful benchmark for understanding biological systems.

Several pieces of previous work have considered optimal strain sensing strategies in insect wings [21, 59, 60, 61]. Although using different sensing tasks and optimization criteria, taken together, they show that remarkably few sensors are needed to detect or identify patterns of strain on the wing. However, this work makes use of sensors that either directly sense strain or encode strain in a continuous variable, unlike the current work which uses spiking sensors. Spiking sensors are not only most faithful to the corresponding biological systems, where primary sensory neurons on the wing encode information in all-or-none action potentials, but they also ensure that sensors transmit temporally sparse signals. Investigating spiking sensors could present opportunities for more efficient, bio-inspired sensing approaches in engineered systems as well.

Although we varied neural encoding properties (nonlinear threshold and filter frequency, Fig 3, Supp Fig 2), all sensors on a given wing have identical encoding properties. Previous work suggests that there are likely two subpopulations of strain-sensitive neurons on the wings of moths [62]. It is possible that a wing with different sensor types may need fewer sensors and adopt a qualitatively different encoding strategy. Additionally, selectivity to multiple features has been demonstrated in neurons in multiple sensory systems, including strain-sensitive neurons in the halteres of crane flies [63, 6]. Allowing individual sensors to respond to multiple stimulus features may also allow for fewer sensors and different sensor positioning. However, it has recently been suggested that all strain-sensitive neurons on insect wings may, in fact, respond similarly in the context of flight, suggesting that sensor placement may be the primary determinant of information encoded by a given sensor [64].

### Sensing needs are one of many evolutionary demands on wing structure

An organism’s evolution reflects a combination of evolutionary constraints and an organism’s needs [65, 66]. For insect wings, these needs may include force generation for flight, sensing, mating display, and predator avoidance, among others [67, 68, 69]. How wing structure impacts aerodynamic performance has been extensively studied [15, 16, 17, 18, 20, 70, 71, 72]. These studies generally point to the importance of stiffness gradients (from base to tip and from leading edge to trailing edge) and rigidity provided by wing veins. However, the consequences of wing structure for sensing have remained unexplored, despite the fact that insect wings have long been known to provide sensory feedback during flight [73, 10, 74].

Incorporating additional constraints may shed light on how these needs interact. For example, in insects, campaniform sensilla on the wings are restricted to be on or near wing veins [75, 62, 76, 77]. Because we used a highly simplified wing model that did not include wing veins (or any spatial variation in stiffness), we allowed sensors to be placed at any point on a dense grid over the surface of the wing. Constraining sensor locations is straightforward and can be achieved by incorporating an additional penalty in the optimization step to decrease the likelihood of sensors being located in particular regions.

In the present work, we focus on how wing structure affects sensing and do not explore how this might interact with other roles of the wing, such as actuation, predator avoidance, and mating display. An important, and challenging, open question is: how are these diverse needs balanced over evolutionary timescales? Answering this question demands an interdisciplinary approach that combines biomechanics, neuroscience, and evolutionary biology to produce a more complete understanding of the processes driving wing evolution.

## Methods

Code that implements strain simulations, conversion of strain to spiking data, and sensor optimization can be found at https://github.com/aiweber/optimal_sensing_ELwing.

### Euler-Lagrange simulations of wing strain

We first simulate spatiotemporal patterns of strain over the surface of the wing using a previously developed Euler-Lagrange model based on parameters of wings of the hawkmoth *Manduca sexta* [29]. In this model, the wing is represented by a flat plate with a span of 50 mm, chord length of 25 mm, and thickness of 0.0127 mm. The wing flaps at 25 Hz and is subject to rotation in different axes of comparable magnitude to rotations experienced by a hawkmoth during free flight (constant at 10 rad/s unless otherwise noted) [78].

We vary Young’s modulus to explore the effects of changing structural properties of the wing, namely wing stiffness. The range of explored stiffness values is approximately centered at 3 GPa (stiffness factor =, corresponding to a spanwise flexural stiffness of ~1.5 * 10^*−*4^ Nm^2^, comparable to experimentally measured spanwise flexural wing stiffness in *Manduca* [26]. We vary Young’s modulus values from 0.7 GPa to 10.0 GPa. Below 0.7 GPa, simulations become unstable.

We simulate strain over a dense grid representing all possible sensor locations (1 mm spacing, resulting in 26 chordwise and 51 spanwise sensors, for a total of 1,326 sensor locations). Unless otherwise noted, we use normal strain in the spanwise direction. See [29] for additional details of Euler-Lagrange simulations and [21] for details of simulations with disturbances. We typically simulate 3 seconds of data (75 wingbeats) for each condition at a sampling rate of 10 kHz.

### Strain encoding with neural-inspired spiking sensors

Neural-inspired sensors encode strain following a procedure similar to earlier work [21]. At each sensor location, strain is first convolved with a linear filter, representing the temporal feature of strain which the sensor is most sensitive to. The filtered strain signal reflects the similarity between linear filter (i.e., feature) and the strain experienced by the sensor over time. Filtered strain is then transformed by a static nonlinearity, which determines the sensor’s selectivity for that particular feature: sensors with high threshold will only respond when the temporal pattern of strain is very similar to the feature given by the linear filter, corresponding to strong selectivity. The output of the linear-nonlinear encoding represents a probability that the neuron will generate a spike.

The shapes of the linear filter and nonlinearity are based on previous electrophysiological recordings of responses in mechanosensors of the wing nerve [12]. The linear filter *f* is defined as a decaying sinusoidal function:

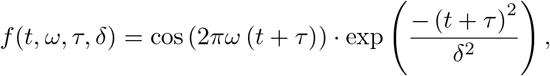

where *ω* is the frequency of the filter, *τ* is the time delay to the peak, and *δ* is the decay time. In this work, 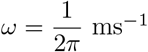, *τ* = 5 ms, and *δ* = 4 ms. The static nonlinearity (also called the nonlinear decision function or nonlinear activation function) is given by:

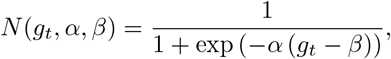

where *g_t_* is the filtered stimulus at time *t*, *α* is the slope parameter, and *β* is the threshold parameter, where the function reaches half-maximum. We hold *α* constant at 50 and vary the threshold *β*. As in previous work, *g* is normalized by a constant (identical for all sensors) such that the maximum value of *g* over all sensors is approximately 1 for wing stiffness of 3 GPa. Threshold is varied between 0.05 and 0.7. (The normalization constant is only important insofar as it sets the scale of the threshold parameter. The range of thresholds we test therefore represent about 5-70% of the maximum filtered stimulus.)

We then generate spikes probabilistically from the output of the linear-nonlinear encoding. The sensor spikes if the probability of firing exceeds a random draw from a standard uniform distribution. We manually impose an absolute refractory period of 15 ms between spikes. This is not intended to represent the actual absolute refractory period of mechanosensors, but rather to empirically match observations from previous experimental work that each sensor fires only 1-2 spikes per wingbeat [79, 73]. For each dataset simulated from the Euler-Lagrange model, we generate 10 sets of spiking responses, for a total of 1,500 data points per optimization (75 wingbeats per Euler-Lagrange simulation, 10 sets of spike generated for each simulation, for both flapping only and flapping with rotation). Noise in the spike generation step dominates noise in the Euler-Lagrange simulations, so similar results would be expected for shorter Euler-Lagrange simulations and more repetitions of spiking responses generated for each simulation.

### Sensor optimization

Our objective in sensor optimization is to determine the placement of a small number of neural-inspired sensors which can be used to determine whether or not the animal is rotating. We simulate spiking data as described above for two cases: one where a wing is flapping, and one where a wing is flapping and rotating (in either the yaw or pitch axis). For both of these cases, we determine the time to first spike within each wingbeat with 0.1 ms precision and use only this information to classify the data. For wingbeats where no spike is elicited, we designate the spike time as zero, though results are unchanged if we instead designate the spike time as a time longer than the wingbeat duration (e.g., 400 ms spike time compared to 40 ms period; see Supporting Information). Data are standardized in this optimization step, but the original (non-standardized) data are used to evaluate accuracy.

To determine optimal sensor locations, we use a previously developed method called *sparse sensor placement optimization for classification* (SSPOC) [80]. This method first uses dimensionality reduction (principal component analysis, in our case) to find a lower-dimensional subspace that captures important features of the data. We then use a linear discriminant analysis (LDA) to find the projection vector *w* that maximally separates the two classes of our data (flapping only, flapping with rotation) in this subspace.

Finally, we use elastic net regularization to solve for a sparse set of sensors *s* that can reconstruct the projection vector *w*:

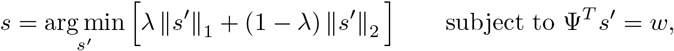

where *s* is a vector of sensor weights (*n*×1, with many near-zero entries), Ψ is the low-dimensional basis (*n*×*m*, *m < n*), *w* is the projection vector in the low-dimensional subspace (*m*×1), and *λ* determines the balance between *L*_1_ and *L*_2_ regularization. We set *m* = 3 and *λ* = 0.9. We use the cvx package to solve this optimization problem (http://cvxr.com/cvx/).

### Performance evaluation

90% of the data is used as a training set for the optimization, and 10% is held out as test data to evaluate accuracy. For straightforward comparison across conditions, we consistently use the top 10 sensors (i.e., sensors with the largest weights in *s*) to assess classification accuracy. LDA is again used to find the best projection vector *w_c_* for the non-standardized test data for only the top 10 sensors. A decision boundary is drawn at the mean of the two condition centroids. (Note that the results will be the same whether or not the test dataset is standardized. We choose to calculate accuracy based on the non-standardized data to highlight the fact that a linear decoding scheme can be used to read out information from the original spike-timing data.)

## Acknowledgments

We would like to thank Thomas Mohren and Michelle Hickner for helpful discussions that shaped this project and for contributions to code development. This work was supported by the Washington Research Foundation (AIW), the eScience Institute at the University of Washington (AIW), and the Air Force Office of Scientific Research MURI FA9550-19-1-0386 (BWB,TLD).

## Supporting information

### Classifying rotation with chordwise strain

Performance is far lower if classification is based on chordwise strain rather than spanwise strain. For classifying rotation in the yaw direction (0 vs 10 rad/s), performance does not rise above chance. Results for classifying rotation in the pitch direction show a small drop in performance and, interestingly, a shift of optimal sensor locations from base to tip compared to classification with spanwise strain (Supp Fig 1).

**Supplementary Figure 1.**
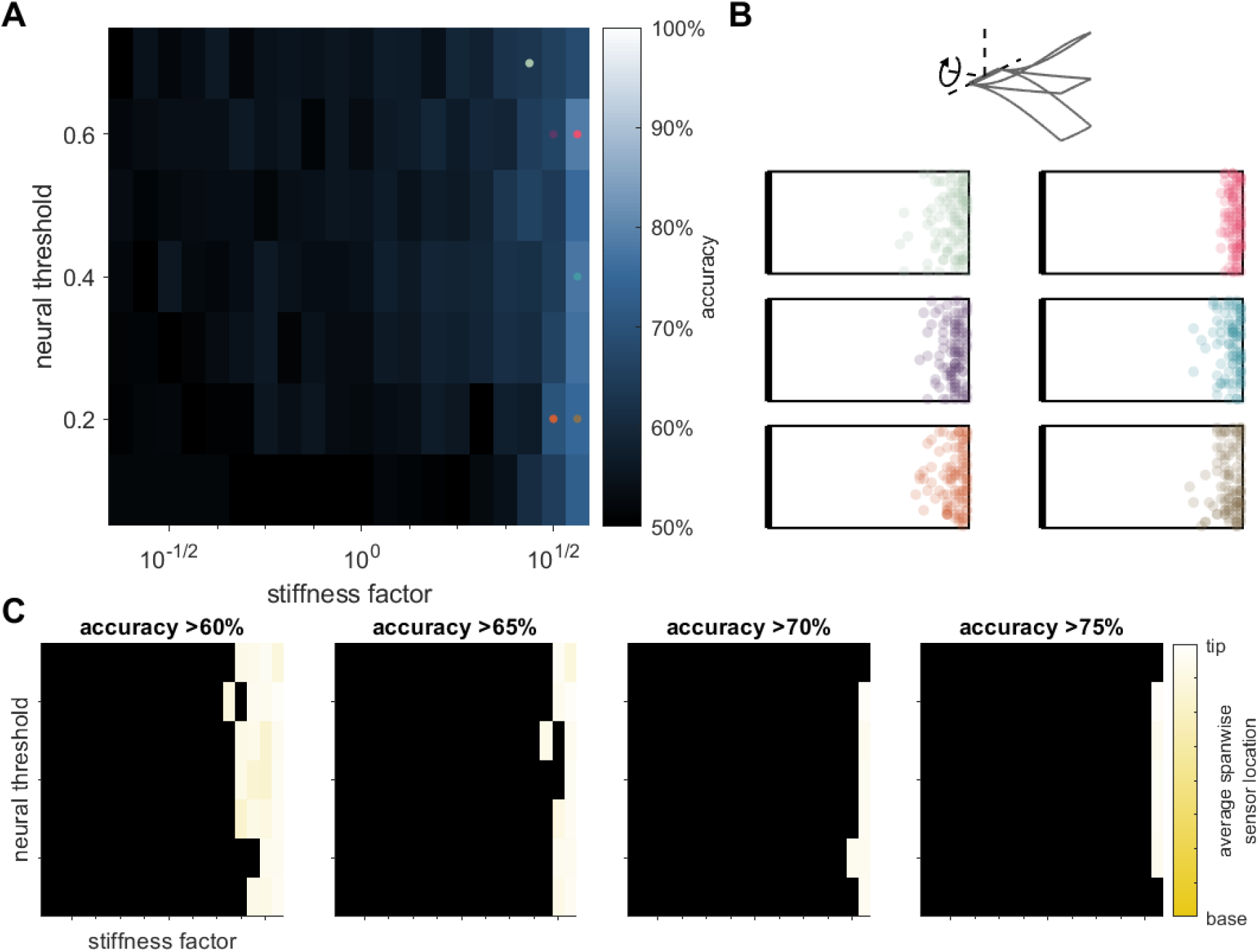
Detecting rotation with chordwise strain reduces accuracy and changes locations of optimal sensors. A: Accuracy as a function of neural threshold and wing stiffness. B: Optimal sensor locations of 10 best sensors, overlaid for 20 different datasets. Neural threshold and wing stiffness of each panel are indicated by the corresponding colored dot in A. Top schematic indicates the axis of rotation. C: Optimal sensor locations in the spanwise direction from wing base (yellow) to wing tip (white) as a function of neural threshold and wing stiffness. Each panel shows results for a different accuracy cutoff. Black indicates parameter combinations that fall below the cutoff. Note the overall lower accuracy and change in optimal sensor locations compared to Fig 4, where spanwise strain is used for classification.

### Effects of changing the frequency of the linear filter

The filter frequency in this work is chosen to correspond to previous work [21], which in turn reflects experimentally measured linear filters [12]. We assess the effects of changing the filter frequency while holding other parameters of neural encoding constant. Nonlinear threshold is held at 0.1, and all other parameters are as indicated in the Methods. Filters of different frequencies are normalized by the sum of squared values at all time points.

Interestingly, classification accuracy depends most strongly on stiffness for this value reflecting experimental observations (indicated by gray box in Supp Fig 2, filter frequency parameter = 1, corresponding to *ω* = 1*/*2*π* ms^*−*1^) compared to higher and lower frequency filters. For higher frequency filters, performance generally does not rise above chance, due to the fact that strain signals contain little to no power at these frequencies. At lower frequencies and when the filter is entirely flat (performing an average over the length of the filter) classification accuracy is nearly 100% across the range of wing stiffness values tested. The experimentally observed frequency content of linear filters in mechanosensory neurons therefore appears to interact particularly strongly with other features of the system (stiffness and neural threshold).

**Supplementary Figure 2.**
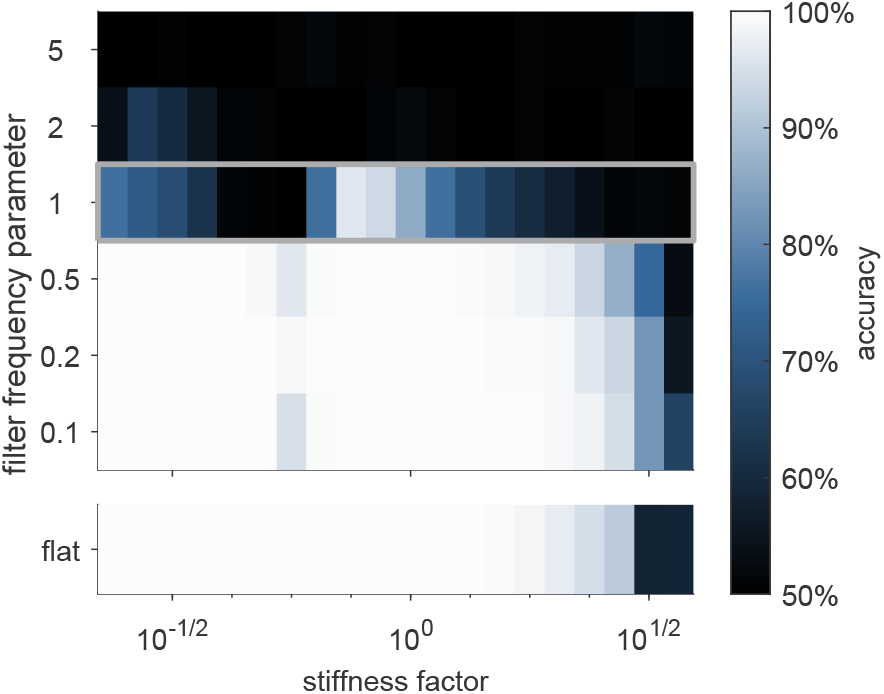
Effects of changing linear filter frequency. Accuracy in detecting rotation in the yaw axis as a function of filter frequency and wing stiffness. Gray box indicates filter frequency used in the rest of the paper. Neural threshold is held constant at 0.1.

### Performance evaluation with a nonlinear classifier

To determine the effects of our choice of linear classifier on our results, we also test the effects of using a nonlinear classifier to determine accuracy. As before, LDA is used to find the best projection vector *w_c_* for stiffness factor the non-standardized test data for only the top 10 sensors. Boundaries are then drawn based on the Gaussian approximations of each category projected onto *w_c_*. Generally only one boundary is used, corresponding to the intersection of the two Gaussian approximations, though in cases where one category has zero variance or variance is both categories is large, resulting in two intersections of the Gaussian distributions, two boundaries are used. We find that this results in only negligible changes to the overall accuracy (Supp Fig 3). Note that sensor locations remain identical to those in Fig 3, as only the classification scheme has changed, while the method of finding optimal sensor locations remains the same as before.

**Supplementary Figure 3.**
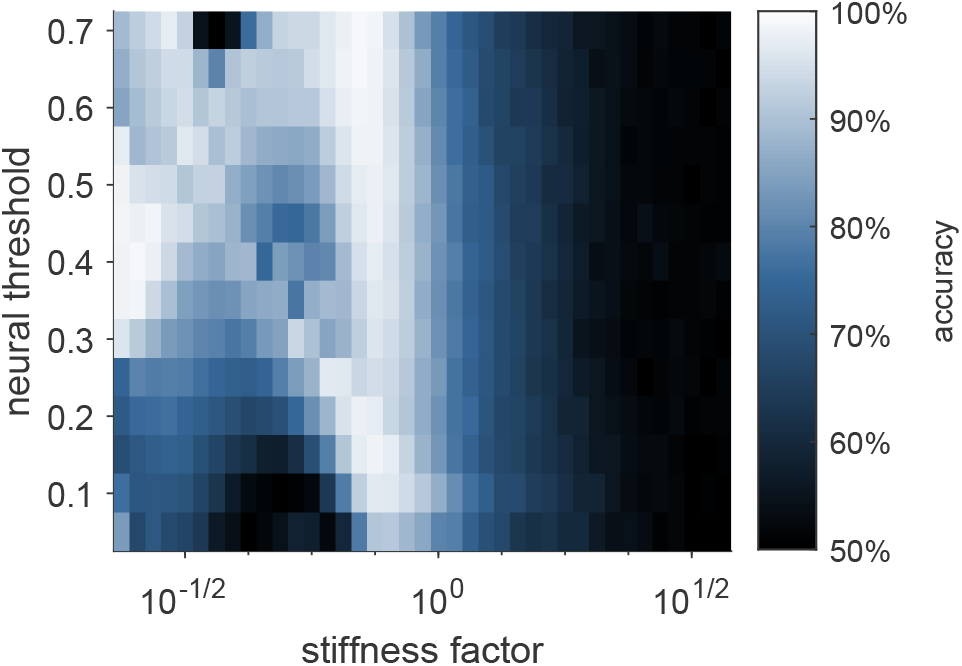
Accuracy of detection of rotation in the yaw axis with a nonlinear classifier. Task accuracy is indistinguishable from a linear classifier.

### Effects of changing the phase of the wingstroke that is defined as time 0

Because classification is based on the time to first spike within each wingstroke, it is possible that the phase of the wingstroke that is defined as t=0 may affect our results. In the above work, time zero is roughly the time of the greatest increase in wing strain over the course of the wingstroke. To test the impact of this, we shift the time defined as time zero by one half wingstroke (20 ms).

The results of this change have small effects on overall accuracy and optimal sensor locations (Supp Fig 4). Though there are slight differences in sensor locations for different neural threshold/stiffness factor combinations, the same distinct regions remain with sensors concentrated at either the wing base corners or edges near or at the wing tip.

**Supplementary Figure 4.**
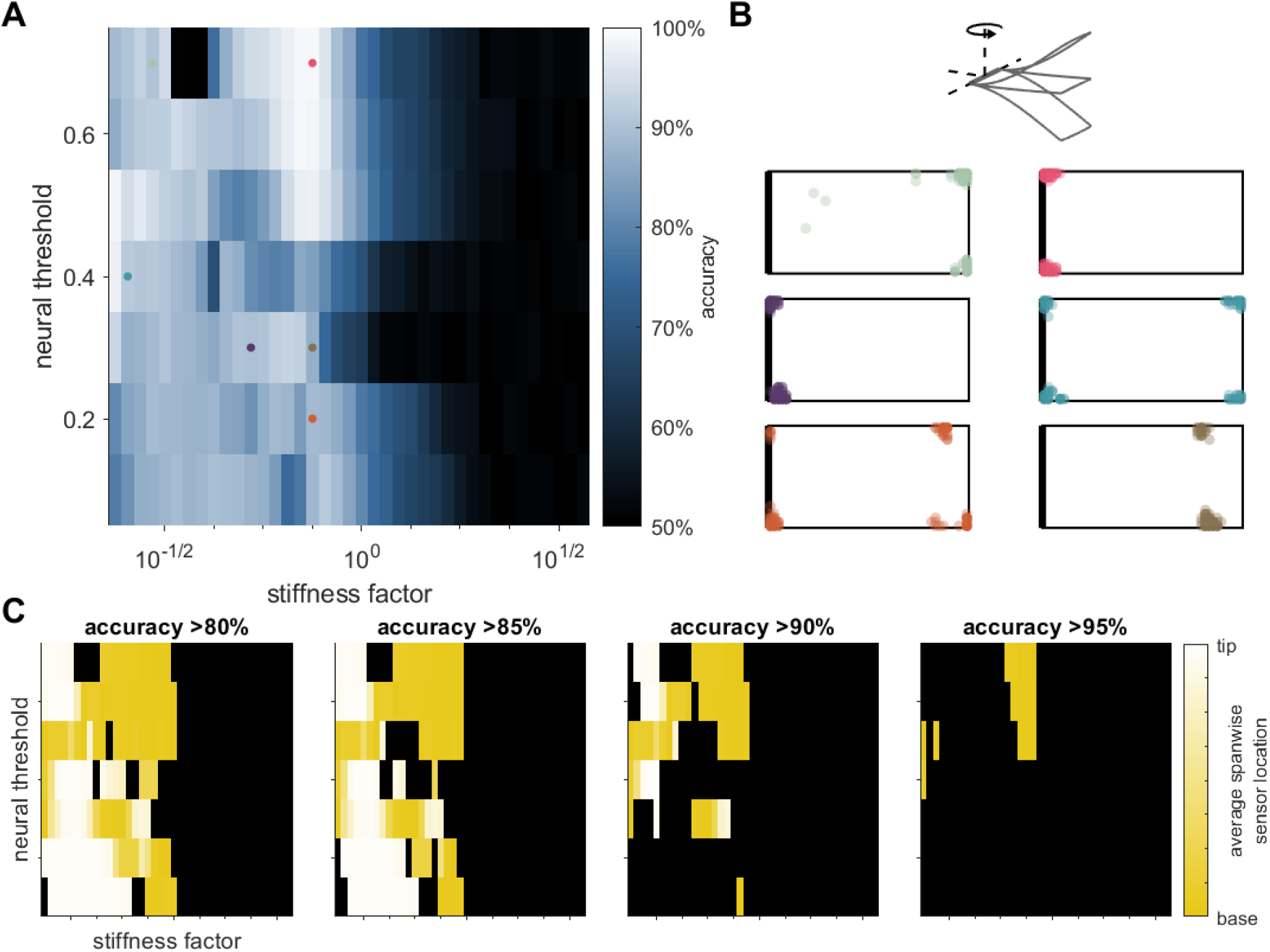
Effects of changing the defined beginning of the wingstroke by 20 ms. A: Accuracy as a function of neural threshold and wing stiffness. B: Optimal sensor locations of 10 best sensors, overlaid for 20 different datasets. Neural threshold and wing stiffness of each panel are indicated by the corresponding colored dot in A. Top schematic indicates the axis of rotation. C: Optimal sensor locations in the spanwise direction from wing base (yellow) to wing tip (white) as a function of neural threshold and wing stiffness. Each panel shows results for a different accuracy cutoff. Black indicates parameter combinations that fall below the cutoff.

### Simultaneous classification of yaw and pitch rotation

In the preceding work, we show results for detection of rotation in the yaw and pitch axes separately. We wished to determine if using a single set of sensors to classify possible rotation in either of these axes would produce unexpected results compared to classifying rotation in each of these axes with a different set of sensors. We perform a separate set of analyses in which a single set of sensors was optimized for 3-way classification of: (1) flapping only, (2) rotation in yaw, and (3) rotation in pitch. Sensor optimization is conducted following the above procedures except that the minimization is subject to Ψ^T^ *s′ w ∊* (rather than Ψ^T^ *s′* = *w*), where *∊* = 10^*−*6^. Once sensor locations are found, we evaluate accuracy by again using LDA to find the best projection vector *w_c_* for the non-standardized test data for only the top 10 sensors.

We then find the centroid of each class projected onto *w_c_* and classify individual points according to the nearest centroid.

Rather than showing fundamentally different trends, classification accuracy and sensor locations reflect a mixture of the two individual classification tasks (Supp Fig 5). Accuracy fails to rise about 70% for any combination of wing stiffness and neural threshold (Supp Fig 5A,C), reflecting a failure to accurately classify all three conditions. Sensor locations reflect a combination of high-concentration in the corners – as observed for yaw – and dispersed locations near the wing base – as observed for pitch (Supp Fig 5B).

**Supplementary Figure 5.**
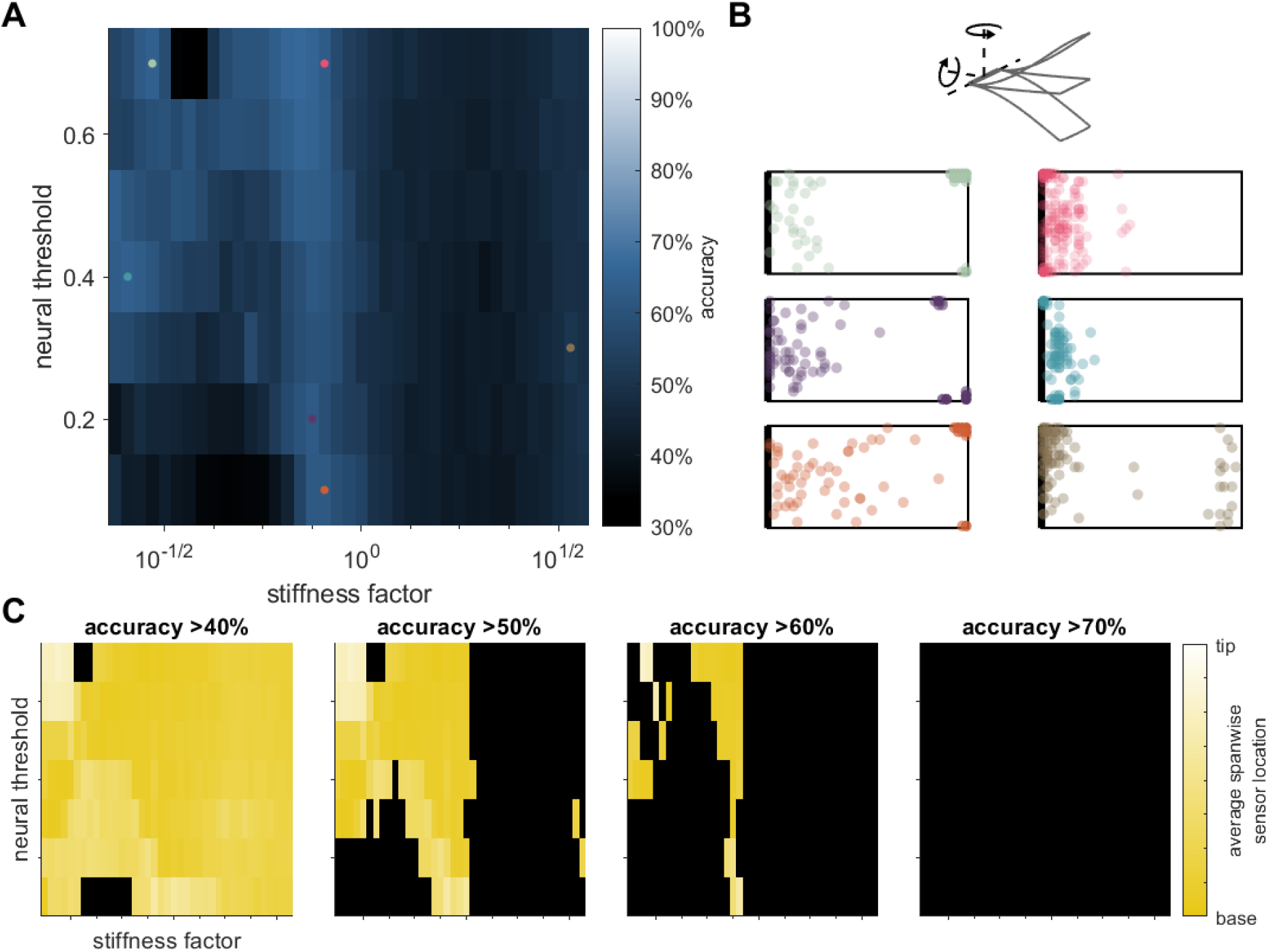
3-way classification of flapping only, rotation in yaw, and rotation in pitch. A: Accuracy as a function of neural threshold and wing stiffness. Note that chance performance is 33%. B: Optimal sensor locations of 10 best sensors, overlaid for 20 different datasets. Neural threshold and wing stiffness of each panel are indicated by the corresponding colored dot in A. Top schematic indicates the possible axes of rotation. C: Optimal sensor locations in the spanwise direction from wing base (yellow) to wing tip (white) as a function of neural threshold and wing stiffness. Each panel shows results for a different accuracy cutoff. Black indicates parameter combinations that fall below the cutoff.

